# A Novel Gene *ARHGAP44* for Longitudinal Changes in Glycated Hemoglobin (HbA1c) in Subjects without Type 2 Diabetes: Evidence from the Long Life Family Study (LLFS) and the Framingham Offspring Study (FOS)

**DOI:** 10.1101/2024.05.16.594575

**Authors:** Siyu Wang, Petra Lenzini, Bharat Thygarajan, Joseph H. Lee, Badri N. Vardarajan, Anatoli Yashin, Iva Miljkovic, E. Warwick Daw, Shiow J. Lin, Gary Patti, Michael Brent, Joseph M. Zmuda, Thomas T. Perls, Kaare Christensen, Michael A. Province, Ping An

## Abstract

Glycated hemoglobin (HbA1c) indicates average glucose levels over three months and is associated with insulin resistance and type 2 diabetes (T2D). Longitudinal changes in HbA1c (ΔHbA1c) are also associated with aging processes, cognitive performance, and mortality. We analyzed ΔHbA1c in 1,886 non-diabetic Europeans from the Long Life Family Study to uncover gene variants influencing ΔHbA1c. Using growth curve modeling adjusted for multiple covariates, we derived ΔHbA1c and conducted linkage-guided sequence analysis. Our genome-wide linkage scan identified a significant locus on *17p12*. In-depth analysis of this locus revealed a variant rs56340929 (explaining 27% of the linkage peak) in the *ARHGAP44* gene that was significantly associated with ΔHbA1c. RNA transcription of *ARHGAP44* was associated with ΔHbA1c. The Framingham Offspring Study data further supported these findings on the gene level. Together, we found a novel gene *ARHGAP44* for ΔHbA1c in family members without T2D. Follow-up studies using longitudinal omics data in large independent cohorts are warranted.

## Introduction

Glycated hemoglobin (HbA1c) is utilized both for the diagnosis and monitoring of type 2 diabetes (T2D), indicative of glycemic control and long-term complication risks in T2D management (Sherwani et al., 2016). HbA1c levels have a genetic basis. More than 120 loci associated with HbA1c have been identified in individuals without T2D through genome-wide association studies (GWAS) (Chen et al., 2021). Linkage scans have also revealed significant genomic regions influencing HbA1c (Meigs et al., 2002, 2007). Some of those regions and genes have been confirmed in multi-ancestry cohorts (Sarnowski et al., 2019).

Despite the acknowledged importance of HbA1c for T2D diagnosis and management, there remains a lack of longitudinal studies focusing on the long-term changes in HbA1c levels (ΔHbA1c). Most existing research tends to focus on the short-term fluctuations and control of HbA1c in T2D patients. However, understanding the long-term trends and changes in HbA1c levels, especially in populations without T2D, is crucial for developing more effective strategies for pre-diabetes diagnosis and improving healthy aging.

A previous GWAS from the Long Life Family Study (LLFS) confirmed two known common loci at *GCK* and *HK1* and uncovered 25 suggestive loci for influencing baseline HbA1c among non-diabetic participants of the LLFS (An et al., 2014). In the present study, we conducted a GWAS of ΔHbA1c followed by a linkage-guided sequence analysis under a significant linkage peak on *17p12* using the latest available whole genome sequencing and omics data, and further pursued replication using the Framingham Offspring Study (FOS) data.

## Design and Methods

### Cohort Populations

The LLFS is a comprehensive, international, longitudinal study that spans two European ancestry generations, focusing on longevity and the factors underlying healthy aging (Wojczynski et al., 2022). The LLFS was conducted at four field centers, three in the United States (Boston, Pittsburgh, New York) and one in Denmark. It enrolled 4,953 participants from 539 families between 2006 and 2009. The visit 1 collected data on anthropometrics, blood pressure, physical performance, pulmonary function, and various blood tests. A second visit from 2014 to 2017 replicated the initial protocols and added carotid ultrasonography measures. HbA1c levels were measured at the University of Minnesota’s Advanced Research and Diagnostics Laboratory using high-performance liquid chromatography (HPLC). The measurements employed Tosoh analyzers calibrated to the National Glycohemoglobin Standardization Program’s standards. The laboratory’s precision for HbA1c values showed a coefficient of variation ranging from 1.4% to 1.9%. ΔHbA1c, derived by growth curve modeling using HbA1c collected from two exams seven years apart, was adjusted for age, sex, BMI, smoking, field centers and 20 principal components, and blom-transformed to approximate normality prior to genetic testing. Subjects with clinical diagnosis of T2D or T2D treatment and undiagnosed T2D cases whose fasting glucose ≥ 126 mg/dl or HbA1c ≥ 6.5% were excluded from this analysis.

### Sequencing Data Preparation

Genotyping was performed on participants using the Illumina Human Omni 2.5 v1 chip by the Center for Inherited Disease Research (CIDR), leading to 1,421,289 SNPs after applying quality controls for call rate<98%, minor allele frequency (MAF) < 1%, p value Hardy–Weinberg equilibrium < 1e−6, and correct correspondence with the 1000 Genomes Project.

Whole genome sequencing (WGS) was executed using Illumina platforms at the McDonnell Genome Institute (MGI), Washington University, with reads aligned to GRCh38. Variant calling followed a four-step process using GATK tools, with additional QC to eliminate contaminated samples and those with hunsuitable coverage or high Mendelian errors.

The visit 1 RNA sequencing was performed on extracted RNA from PAXgene™ Blood RNA tubes using the PAXgene microRNA extraction kit. Library preparation and quality control was managed by the Division of Computation & Data Sciences, Washington University. The nf-core/rnaseq 3.14.0 pipeline facilitated read alignment, duplication marking, and transcript quantification. Post-processing included filtering out samples with a high fraction of reads mapping to intergenic regions, filtering out genes with very low expression levels, normalizing gene counts with variance stabilizing transformation in DESeq2, leading to a final selection of 1,810 samples and 16,418 genes.

Lipid metabolomic profiling at visit 1 was executed at Washington University’s Biomedical Mass Spectrometry Lab. The laboratory implemented LC/MS for untargeted lipid detection, matched polar metabolites with internal and online databases, and annotated MS/MS lipid data. After rigorous QC and batch effect correction using a pooled QC sample, the analysis yielded data on 188 lipids across 13 compound classes.

### Framingham Heart/Offspring Study for Replication

The Framingham Heart/Offspring Study (FHS/FOS) is a longitudinal cohort study tracking three generations for up to 65 years to assess cardiovascular disease risk factors. HbA1c measurements were taken from exam 5 and exam 7, selected for their similarity with LLFS intervals. Longitudinal HbA1c employed the same growth curve modeling adjusted for age and sex. Its genomic research, including whole genome sequencing (WGS) and related phenotypic analysis, is incorporated into the NHLBI’s Trans-Omics for Precision Medicine (TOPMed) Whole Genome Sequencing Program. WGS (freeze 9b) data from the Framingham Offspring Generation was used for replication. Bi-allelic single nucleotide variants (SNVs) that pass all QC filters were kept, resulting in ∼52 million variants. Samples were excluded if they were either sequence controls, not sequenced in blood, or had FREEMIX percentage > 3%. Additionally, samples were removed if they had a mean depth of < 30x or < 95% of sites covered at 10x or < 80% at 20x. A total of 2,186 participants, derived from the offspring cohort’s sequencing data, were queried for replication.

### Statistical Analysis

The Sequential Oligogenic Linkage Analysis Routines (SOLAR) program is designed to accommodate familial relatedness by employing maximum-likelihood based methods. These methods are utilized to estimate the residual genetic heritability of outcome measures, as well as to discern the variance that can be attributed to fixed covariate effects (Almasy & Blangero, 1998). We used SOLAR to select linked families with PEDLOD > 0.1 or PEDLOD/N > 0.01, where N was family members included in the pedigree trait count, are defined as the “potentially linked” of families. Then we ranked the families by LOD and the minimum number of families with the highest LODs that add up to at least 9 were defined as the “top linked” families. We performed GWAS analyses using a linear mixed model for additive dosage of the variants. Familial relationships were accounted for by including a kinship matrix, estimated via the “kinship” R package, as a random effect in the “lmekin” R package. LODs > 3 for GWLS and *p* < 5e-8 for GWAS association were used to declare significance. Associations between phenotype and RNASeq or phenotype and metabolites were analyzed using similar linear models adjusted for age, sex, and field centers.

## RESULTS

### Demographic Characteristics

This analysis included a total of 1,886 family members (826 men and 1,060 women) from LLFS with complete phenotypic data at both visits and genotypic information (Table 1). Similar inclusions and exclusions were applied in the replication cohorts and a total of 1,739 (752 men and 987 women) from the FOS with complete phenotypic data at both visits and genotypic information were selected for replication (Table 1). Detailed characteristics of the LLFS linkage-enriched group and others are also given in Table 2. We identified 176 subjects from 16 linkage-enriched families (“top linked” families with cumulative LODs over 9) of in the LLFS. Significant mean differences in characteristics were noted both between the study group and the others, as well as within each group across sexes. Additionally, the two cohorts differ significantly in terms of HbA1c changes; the LLFS shows minimal change in HbA1c levels, whereas the FOS exhibits a more substantial change (Table 1).

**Table 1.**
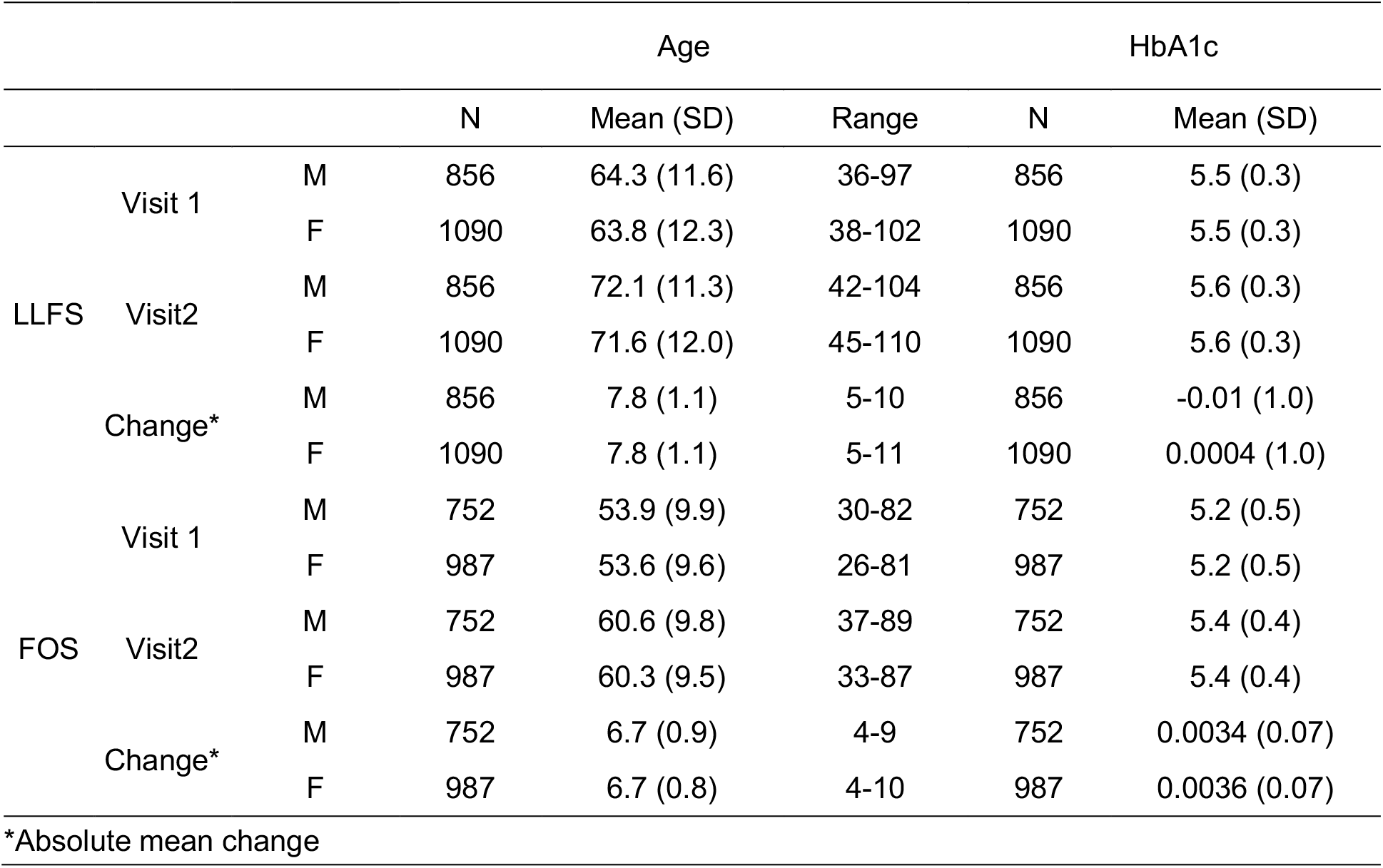
Sample characteristics of the LLFS and FOS cohorts.

**Table 2.** Sample characteristics of the linkage-enriched families in the LLFS.

|  |  | Men |  |  | Women |  |  |
| --- | --- | --- | --- | --- | --- | --- | --- |
|  |  | N | Mean | SD | N | Mean | SD |
| Study Group<br>(N=176) | Age (years) | 78 | 60.89† | 10.64 | 98 | 61.59† | 12.05 |
|  | Fasting Glucose (mg/dL) | 78 | 96.19*† | 9.68 | 98 | 92.22*† | 8.15 |
|  | HbA1c (%) | 78 | 5.42 | 0.34 | 98 | 5.38† | 0.3 |
|  | BMI (kg/m<sup>2</sup>) | 77 | 28.16*† | 3.35 | 98 | 26.44* | 4.81 |
|  | ΔHbA1c | 78 | -0.04 | 1.13 | 98 | -0.05 | 1.13 |
|  | HOMA-B | 70 | -0.04 | 0.92 | 93 | 0.01 | 0.89 |
|  | HOMA-IR | 70 | -0.05 | 0.94 | 93 | 0.01 | 0.88 |
|  | TG/HDL Ratio Growth Curve | 57 | 0.04 | 0.97 | 92 | -0.19 | 0.85 |
| Other<br>(N=1710) | Age (years) | 748 | 64.79† | 11.78 | 962 | 64.37† | 12.34 |
|  | Fasting Glucose (mg/dL) | 743 | 93.05*† | 11 | 959 | 89.26*† | 10.4 |
|  | HbA1c (%) | 743 | 5.47 | 0.31 | 962 | 5.47† | 0.3 |
|  | BMI (kg/m<sup>2</sup>) | 728 | 27.16*† | 3.46 | 944 | 26.22* | 4.56 |
|  | ΔHbA1c | 748 | -0.01 | 0.99 | 962 | -0.0009 | 0.96 |
|  | HOMA-B | 674 | -0.02 | 1.02 | 881 | -0.05 | 0.96 |
|  | HOMA-IR | 674 | -0.06 | 1.01 | 881 | -0.09 | 0.98 |
|  | TG/HDL Ratio Growth Curve | 420 | 0.03 | 0.99 | 632 | -0.004 | 1.01 |

\*Significant ( $p < 0.05$ ) mean differences between sexes
†Significant ( $p < 0.05$ ) mean differences (within sex) between subsets
HOMA-B, HOMA-IR: HOMA derived of $\beta$ -cell function, insulin resistance

### Discovery in LLFS

Genome-wide linkage analysis yielded a significant region on *17p12* (Fig. 1A. LODs = 3.59, 13 Mb). Heritability reached 36.8% with a *p*-value of 1.4e-9 (standard error = 6.8%). Whole genome sequencing association analysis did not find any significant variants with the *p* < 5e-8 (Fig. 1A), but it did reveal 19 suggestive variants with 5e-8 < *p* < 1e-5, including one in *HK1* (rs16926246, *p* = 3.05e-7) that was previously reported for baseline HbA1c in the LLFS. The suggestive variant in the *RAP1GAP2* (rs143842515, *p* = 2.51e-7), located 1Mb upstream of the linkage peak, was identified. A variant in the *ARHGAP44* fell in 1-LOD-support interval (Fig.1B rs56340929, *p* = 1.77E-06, MAF=6%, accounting for linkage = 26.5%).

**Figure 1A.**
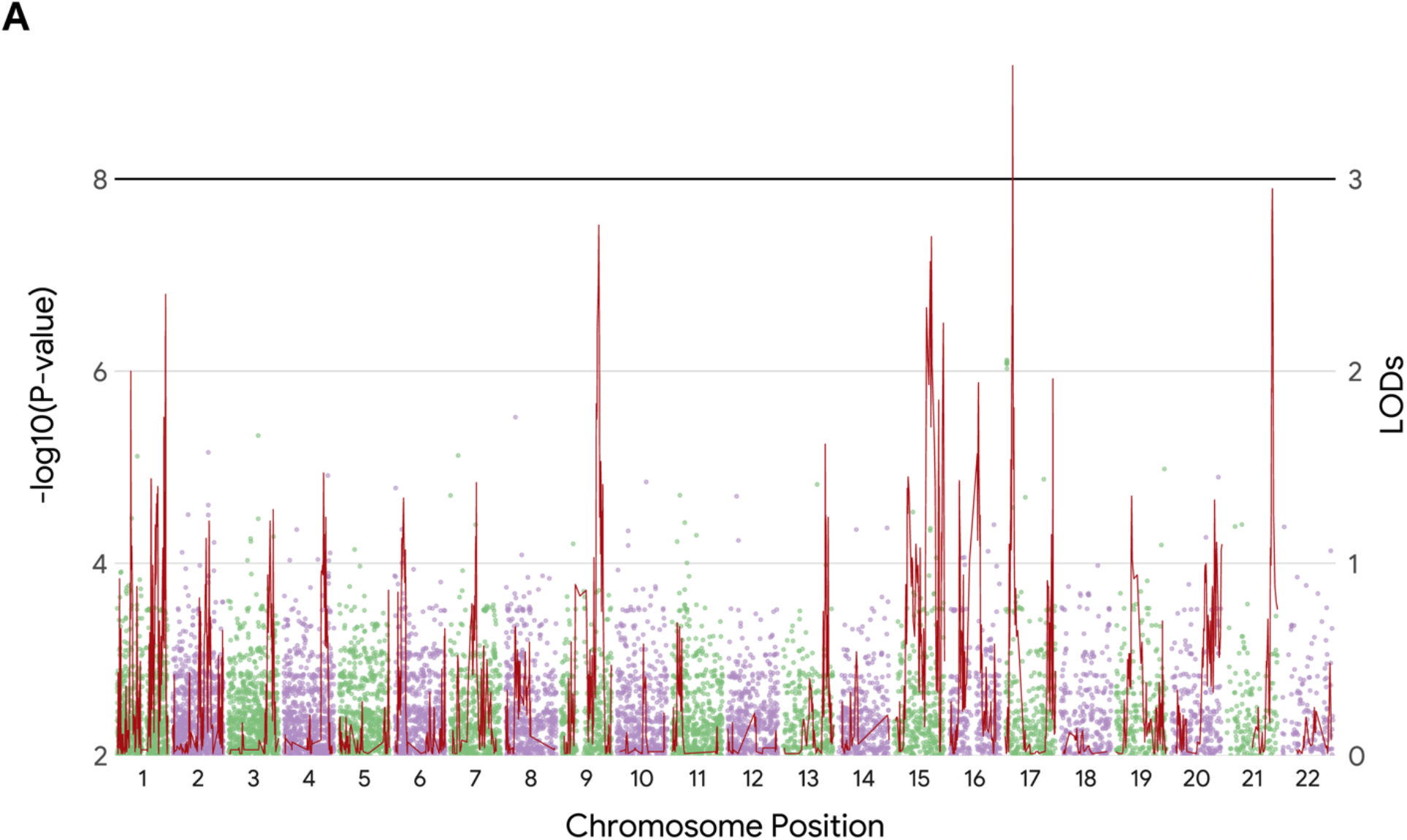
An overlay of Manhattan plot from the GWAS and GWLS results. The illustration synthesizes our findings in the overall data. The horizonal black reference line denotes the established criteria for genome-wide significance (*p* < 5e-8), and linkage scan significance (LODs > 3.0).

**Figure 1B.**
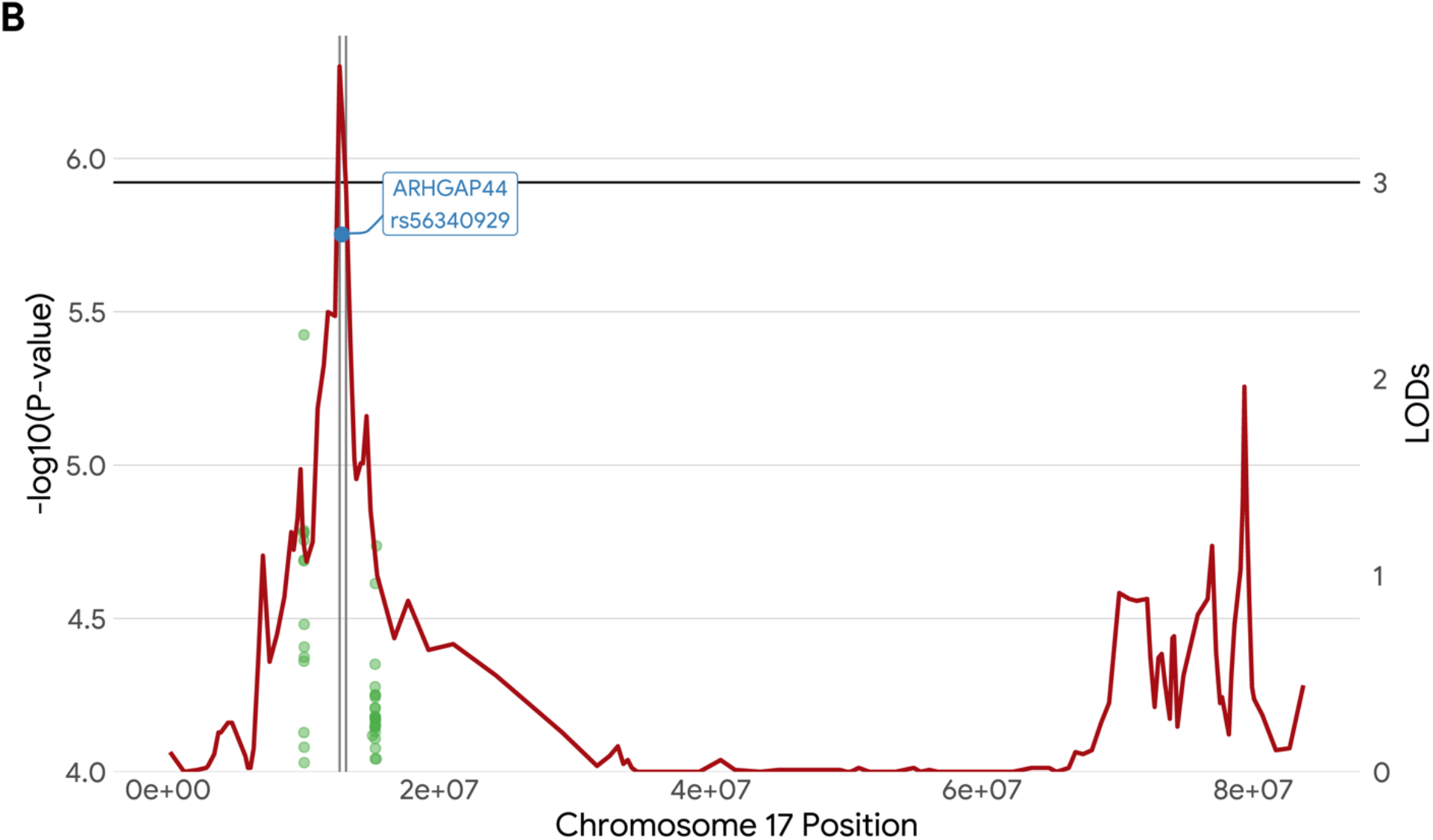
Highlights of the identified lead SNP rs56340929 under the right linkage peak (*17p12*) in the linkage-enriched families. The black line denotes a threshold at LODs > 3.0, while vertical gray lines represent the 1-LOD support interval. SNP rs56340929 is located within this interval.

Taking advantage of our available transcriptome sequencing (RNAseq) data, we assessed the association between quantification of the *ARHGAP44* RNA transcript and ΔHbA1c among 176 subjects, and we found they were significantly associated (*β* = −0.0002, SE = 0.002, *p* = 0.02). We further explored the association of the *ARHGAP44* RNA transcript and its corresponding SNP rs56340929, we found that the association was only marginally missed (*β* = −0.21, SE = 0.16, *p* = 0.07). We assessed currently available metabolomics data (188 metabolites in 13 compound classes) among the 16 linkage-enriched families. We found that triacylglycerol (*p* = 0.024) and sphingomyelin (*p* = 0.036) appeared to be marginally associated with ΔHbA1c. These however were non-significant after the *p*-values were corrected for multiple testing (*p* < 3e-4 for metabolites or *p* < 0.004 for compound classes). We also assessed rs56340929 associations with the currently available lipidomic data; and we found no significant associations after correction for multiplicity. This would not suggest at least from this analysis that the *ARHGAP44* gene is directly engaged in regulating lipid metabolism.

### Replication in the FOS

A total of 1,739 non-diabetic subjects from the FOS with complete phenotypic data at both visits and genotypic information were used for replication. Another SNP rs140270267 (830 bp downstream of the rs56340929, *p* = 0.0002, MAF = 2%) was identified, which indicated that the *ARHGAP44*-rs56340929 was only replicated at a nearby site (rs140270267, D’ = 1, R^2^ = 0.002, not in linkage disequilibrium, not at an exact SNP site) or at the *ARHGAP44* gene level (SimpleM, N_eff_ = 148, *p* = 0.00029 < 0.00034).

## DISCUSSION

This analysis found an appreciable and significant genetic component (heritability of 37%) for ΔHbA1c among non-diabetic subjects from the LLFS. The heritability appears to be compatible with or slightly lower than our previously reported estimate of 42% in the LLFS (An et al., 2014) and estimates of 47-59% in other family studies (Meigs et al., 2002; Pilia et al., 2006; Soranzo, 2011) for HbA1c at baseline. No other familial aggregation reports of ΔHbA1c are noted. Interestingly, this analysis identified a significant linkage peak on *17p12* (LODs = 3.6) for ΔHbA1c. Several studies found suggestive linkage evidence on this region for relevant traits including fasting glucose (Loos et al., 2003), circulating leptin (Kissebah et al., 2000), T2D (Lindgren et al., 2002), coronary artery disease (Gao et al., 2014), and metabolism syndrome (Kissebah et al., 2000; Zhang et al., 2013). While no previous linkage scans were found for ΔHbA1c, Meigs and colleagues reported a suggestive linkage on chromosome 1 for baseline HbA1c in the FHS (Meigs et al., 2002). Interestingly, in this analysis, we found that our linkage peak on *17p12* was substantially attenuated (LODs from 3.6 to 1.0) when ΔHbA1c was corrected for baseline HbA1c, whereas it only modestly changed when ΔHbA1c was corrected for baseline fasting glucose levels (LODs from 3.6 to 3.4) or baseline hemoglobin levels (LODs from 3.6 to 3.8). This observation would suggest that the linkage peak for ΔHbA1c may be partially conditional on baseline HbA1c levels, but it does not clearly distinguish between glycemic and non-glycemic (erythrocytic) pathways.

Here, our best GWAS finding of *HK1*-rs16926246 (*p* = 3e-7) for ΔHbA1c did not reach genome-wide significance (*p* < 5e-8). To our knowledge, there are no previous GWAS reports for ΔHbA1c. For baseline HbA1c, the *HK1* is a confirmed locus in the Women’s Genome Health Study (rs7072268, *r*^2^ = 0.13, MAF = 0.50, Pare et al 2008), the MAGIC consortium (rs16926246, MAF = 0.10, Soranzo et al 2010), and the LLFS (rs17476364, *r*^2^ = 0.62, MAF = 0.10, An 2014). These common SNPs seem to be not in perfect linkage disequilibrium, and likely represent independent *HK1* variants. The *HK1* gene encodes hexokinase 1, the enzyme catalyzing the first step of glycolysis, that regulates glucose metabolism. The HK1 deficiency in erythrocytes causes severe non-spherocytic hemolytic anemia. Previous reports also suggested that the *HK1* locus could influence HbA1c levels via erythrocyte biology (Paré et al., 2008; Soranzo et al., 2010).

The only statistically significant discovery from our linkage-guided sequence analysis was the *ARHGAP44-rs56340929* (*p* = 2e-6, MAF = 6%) for ΔHbA1c. We assessed all sequence elements under the linkage peak on *17p12* (within 1-LOD support interval from 12.7 Mb to 12.9 Mb) and found this lead variant accounted for nearly 30% of the linkage peak. Our RNAseq data showed that *ARHGAP44* expression level was only marginally missed its association with the variant (*p* = 0.07) but significantly associated with ΔHbA1c (*p* = 0.02). Our lipidomics data did not reveal any metabolites that were significantly associated (*p* < 3e-4 after multiple testing correction) with ΔHbA1c or with the variant. This observation would suggest *the ARHGAP44* gene variant does not directly regulate lipid metabolism. The *ARHGAP44* encodes Rho GTPase activating protein 44 which is involved in the control of Rho-type GTPases (Xu et al., 2017). The *ARHGAP44* gene has also been reportedly associated with cardiovascular diseases, serum creatinine and glycemic traits including HbA1c (Dornbos et al., 2022). Finally, we looked up the *ARHGAP44*-rs56340929 in the FOS and found encouraging validation evidence at a neighboring variant (rs14270267, 830 bp downstream, *p* = 0.0002, *r*^2^ = 0.002) and at the gene level (simpleM, *p* < 0.0003, N_eff_ = 148, after correction for multiple testing) though not at the exact SNP site (*p* = 0.8).

Strengths of the current study included its extended family design, relatively large sample size, longitudinal data availability with multiple visits, well-defined phenotypic measures, and moreover availability of multi-omics data. However, there are also few limitations needed to be noted in this analysis. Sample heterogeneity and thus genetic heterogeneity may exist across the LLFS and FOS. This was evidenced by significant mean differences in HbA1c levels and key covariates between the two studies (see Table 1). The exceptional longevity of the LLFS sample compared to the general population may introduce selection bias that potentially rendered somewhat data heterogeneity. Additionally, incomplete access to the FOS data with missing covariate variables may impede the exact replication of phenotyping adjustment methods, which may further diminish the validity of the replication.

In conclusion, this integrated linkage-guided sequence analysis allowed for our identification of a novel gene the *ARHGAP44* for ΔHbA1c in the LLFS with supportive evidence from our omics data as well as encouraging replication data using the FOS. Further independent replications from large cohorts are needed to confirm and extend our findings.

## Author contributions

S.W. researched the data and wrote the manuscript. P.L., B.T., J.H. L., B.V., A.Y., I.M., E.W.D., S.J.L., G.P., M.B., J.Z., T.P., K.C., M.A.P., P.A. either wrote paragraphs or sentences or edited the manuscript. All the authors in the LLFS participated in the weekly priority paper teleconference calls where this manuscript was initiated and revised. All the coauthors reviewed and approved submission of the manuscript to Diabetes. S.W. is the guarantor of this work, and as such, had full access to all the data in the study and takes responsibility for the integrity of the data and the accuracy of the data analysis.

## Statements of assistance

The investigators thank all the LLFS participants and staff for their valuable contributions. We are grateful to LeAnne Kniepkamp for her administrative help and effort. The authors thank Lihua Wang, M.D., M.S. for her assistance in statistical data analysis for this manuscript.

## Prior published abstract of the study

Wang S, Thyagarajan B, Lee J, Zmuda J, Christensen K, Province M, An P. Novel gene variants associated with HbA1c changes over time among non-diabetic subjects in the Long Life Family Study. *Innov Aging*. 2023 Dec 21;7(Suppl 1):1088. doi: 10.1093/geroni/igad104.3496. PMCID: PMC10738390.

